# Aversive Conditioning of Spatial Position Sharpens Neural Population-level Tuning in Visual Cortex and Selectively Reduces Alpha-band Activity

**DOI:** 10.1101/2020.11.14.382960

**Authors:** Wendel M. Friedl, Andreas Keil

## Abstract

Processing capabilities for many low-level visual features are experientially malleable, aiding sighted organisms in adapting to dynamic environments. Explicit instructions to attend a specific visual field location influence retinotopic visuocortical activity, amplifying responses to stimuli appearing at cued spatial positions. It remains undetermined, however, both how such prioritization affects surrounding non-prioritized locations, and if a given retinotopic spatial position can attain enhanced cortical representation through experience rather than instruction. This work examined visuocortical response changes as human observers learned, through differential classical conditioning, to associate specific on-screen locations with aversive outcomes. Using dense-array EEG and pupillometry, we tested the pre-registered hypotheses of either sharpening or generalization around an aversively associated location following a single conditioning session. Specifically, competing hypotheses tested if mean response changes would take the form of a gaussian (generalization) or difference-of-gaussian (sharpening) distribution over spatial positions, peaking at the viewing location paired with a noxious noise. Occipital 15 Hz steady-state visual evoked potential (ssVEP) responses were selectively heightened when viewing aversively paired locations and displayed a non-linear, difference-of-gaussian profile across neighboring locations, consistent with suppressive surround modulation of non-prioritized positions. Measures of alpha band (8 – 12.8 Hz) activity and pupil diameter also exhibited selectively heightened responses to noise-paired locations but did not evince any difference across the non-paired locations. These results indicate that visuocortical spatial representations are sharpened in response to location-specific aversive conditioning, while top-down influences indexed by alpha power reduction exhibit all-or-none modulation.

**Significance Statement:** It is increasingly recognized that early visual cortex is not a static processor of physical features, but is instead constantly shaped by perceptual experience. It remains unclear, however, to what extent the cortical representation of many fundamental features, including visual field location, is malleable by experience. Using EEG and an aversive classical conditioning paradigm, we observed sharpening of visuocortical responses to stimuli appearing at aversively associated locations along with location-selective facilitation of response systems indexed by pupil diameter and EEG alpha power. These findings highlight the experience-dependent flexibility of retinotopic spatial representations in visual cortex, opening avenues towards novel treatment targets in disorders of attention and spatial cognition.

---

Humans excel at distilling the statistical regularities of experienced events into general rules of behavior. While accurately associating harmful events with valid warning cues is important for survival, incorrectly linking benign situational cues to noxious events can be maladaptive, and may contribute to various psychopathologies (Lissek et al., 2014; Dunsmoor & Paz, 2015). Characterizing the neurophysiological processes of associating specific sensory cues with aversive outcomes is necessary to advance understanding of this fundamental learning mechanism.

Mounting evidence (e.g. Weinberger, 2004; Chen, Barnes, & Wilson, 2011; Aizenberg & Geffen, 2013; Kass, Rosenthal, Pottackal, & McGann, 2013) indicates that neural representations within primary sensory cortices respond to learned contingencies as well as to basic physical features. How the retinotopic representation of visual field position is affected by such learning, however, remains unknown. Aversive (Classical, Pavlovian) differential conditioning paradigms, which pair an initially neutral stimulus with an intrinsically aversive outcome, offer a well-established method for investigating learning-induced neural response changes. Aversive conditioning studies have, for example, demonstrated that oriented gratings predicting aversive outcomes (CS+ stimuli) come to prompt selectively potentiated visuocortical responses in non-human primates (Li,Yan, Guo, & Li, 2019) and human observers (Moratti & Keil, 2009; Song & Keil, 2014; McTeague, Gruss, & Keil, 2015). Moreover, responses to stimuli varying in physical similarity to the CS+ along an ordered continuum of a manipulated feature can be used to both characterize the extent of acquired changes in sensory sensitivity (Schechtman, Laufer, & Paz, 2010; Holt et al., 2014) and inform as to underlying neural mechanisms (Ringach, Hawken, & Shapley, 2003; Dunsmoor & Paz, 2015; Onat & Büchel, 2015).

Two response distributions widely used to describe neuronal selectivity along a continuum of features are the gaussian (Onat & Büchel, 2015; Tuominen et al., 2019) and the difference-of-gaussian (DoG; Ringach, Hawken, & Shapley, 2003; Yeonan-Kim & Bertalmío, 2016). Applied to aversive conditioning, a monotonic gaussian pattern peaking at the CS+ could suggest that responses to test stimuli are actively graded according to their similarity to the CS+ (Struyf, Zaman, Vervliet, & Van Diest, 2015; Tuominen et al., 2019). Meanwhile, a non-monotonic DoG gradient, as seen when stimuli most physically similar to the CS+ elicit weaker responses than do those that are more distinct, likely implies that a mechanism of suppression is involved (Shapley, 2004; Isaacson & Scanziani, 2011; Angelucci et al., 2017).

In addition to processing biases stemming from the objective characteristics of an observed stimulus, humans invoke numerous higher-order cognitive processes to facilitate stimulus-response association (Rescorla 1988; Dunsmoor, 2015). Changes in neuronal population-level oscillatory activity in the alpha band (~8-14 Hz) have been linked to activation of the attention network (e.g. Capotosto, Babiloni, Romani, & Corbetta, 2009; Romei, Gross., & Thut, 2010), which exerts strong, top-down effects on sensory processing (reviewed in Fiebelkorn & Kastner, 2020). For instance, increased alpha-band activity within sensory cortices is inversely associated with neuronal firing rate (Haegens et al., 2011; Johnston, Ma, Schaeffer, & Everling, 2019) and overall sensory processing (Foxe & Snyder, 2011).

The present study examined the hypothesis that the representation of spatial position within early visual cortex is malleable via aversive conditioning. Spatial location tuning profiles were obtained with steady-state visual evoked potentials (ssVEPs), which emerge primarily from V1 and higher-order visual cortices (Di Russo et al., 2007). Alpha-band amplitude served as an index of extra-striate, higher-order cortical processes during the associative learning process. Gaze fixation was monitored with eye-tracking to ensure consistent retinotopic mapping. Our pre-registered prediction was that within retinotopic visual cortex, neuronal population-level tuning to a threat location would prompt suppression of location representations proximal to the CS+, resulting in a difference-of-gaussian response gradient. Meanwhile, decreases in alpha-band amplitude were expected to show a gaussian profile peaking at the threat-associated location, consistent with less stimulus specific attentive processing of the threat cue. While not part of our pre-registered hypotheses, pupillary responses are presented as an additional measure of learning-induced physiological change.

## Materials and Methods

Testing the primary signals of interest (alpha power and ssVEP amplitude) against our hypotheses generally proceeded as follows: (a) source estimation, (b) testing the fit of gaussian and DoG patterns (figure 1) of experience-dependent signal change across the entire source-estimated scalp topography, (c) PCA of source-space activity, and (d) testing the effect of conditioning as a function of spatial distance from the CS+ for the PCA-weighted signals.

**Figure 1.**
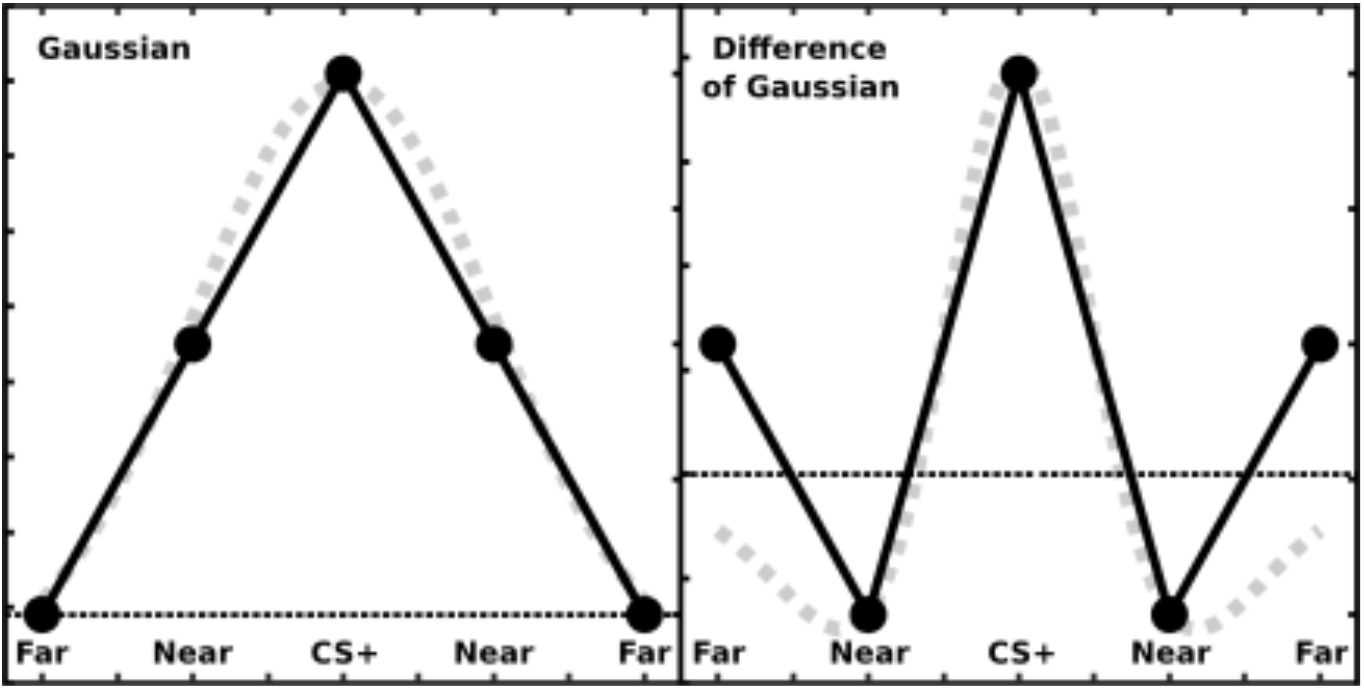
Gaussian and difference-of-gaussian gradients. Grey dotted lines depict each pattern plotted on a continuous horizontal axis. Black lines and black filled circles illustrate the competing hypotheses as tested in the present study with five ordered categorical levels of spatial position. Note that this example fits the hypothesized signal change trend for ssVEP power; these gradients were predicted to be inverted for alpha power change (i.e. maximum alpha decrease at the CS+).

### Participants

Fifty-three students at the University of Florida with normal or corrected to normal vision participated in the study for course credit. All participants reported no personal or family history of epilepsy or photic seizures. One participant chose to discontinue testing due to discomfort with the auditory US, and data from one participant was excluded due to a defective EEG sensor net, leaving a final sample of 51 (19 male; M_age_=19.49, SD_age_=1.22) participants. All participants provided informed consent in accordance with the Declaration of Helsinki and the institutional review board of the University of Florida.

### Stimuli

Visual stimuli consisted of high contrast Gabor patches (1.18 cycles/degree) presented against a black background (0.01 cd/m^2^). Gabor patches subtended 7.07 degrees of visual angle and had a maximum luminance of 96 cd/m^2^. Gabors were presented individually at one of five equally spaced locations, having their inner border at 2.12 degrees of eccentricity from a central fixation dot (figure 2). The small white fixation dot remained on-screen at all times, except during assessment of participants’ awareness of the US / CS+ pairing contingency during acquisition trials (see below). Visual stimuli were presented on a calibrated LCD display (Cambridge Research Systems Display ++) running at 120 Hz connected to a PC running Linux Ubuntu 18.04. Stimuli were created in MATLAB (MathWorks, Natick, MA) with the Psychophysics Toolbox (Brainard, 1997; Kleiner et al., 2007). Gabor patches were flickered at a rate of 15 times per second (Hz) to drive cortical oscillatory responses at the desired steady-state temporal frequency. The unconditioned stimulus (US) was a 90-dB SPL white noise delivered binaurally through two small computer speakers located behind participants’ seated location.

**Figure 2.**
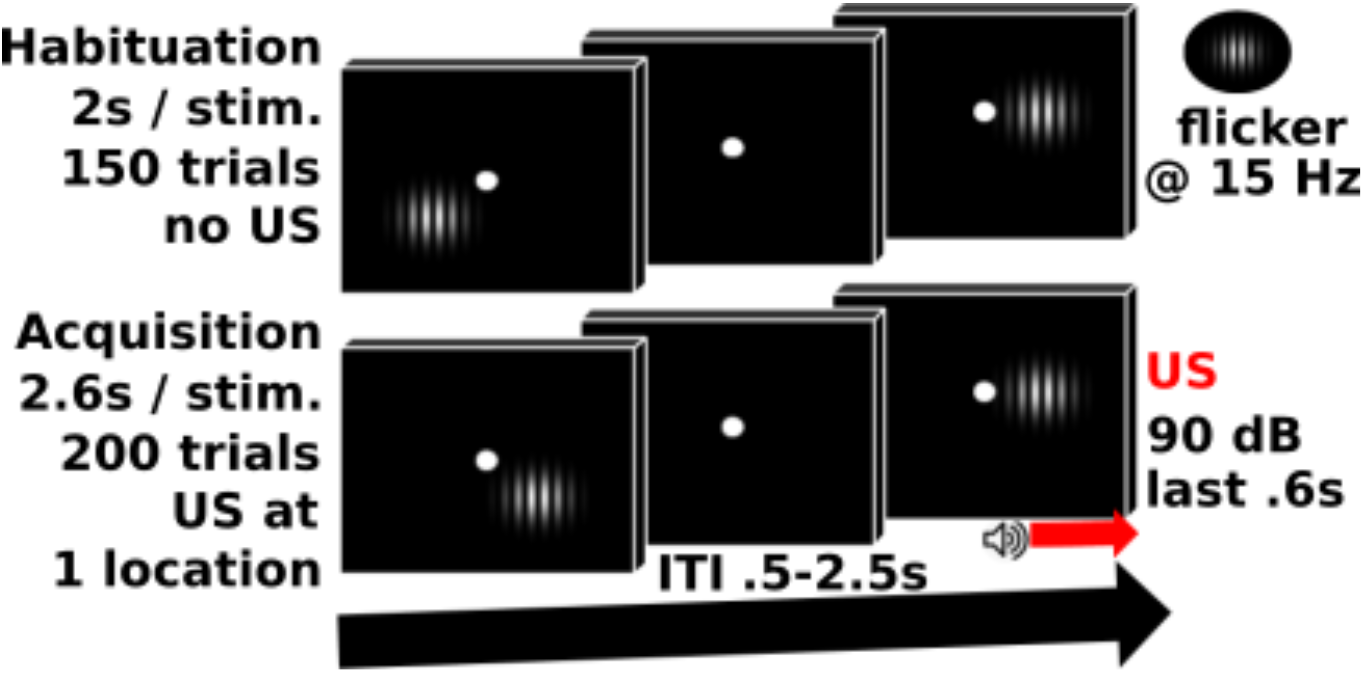
Experimental paradigm. A sequence of two stimulus presentations is shown for habituation (no US noise) and acquisition (90 dB white noise blast at one location) trial blocks.

### Procedure

Participants were seated in a comfortable chair with their heads placed on a chin rest in a dimly lit (∼ 60 cd/m^2^) room. Viewing distance to the display monitor was adjusted to 120 cm and distance to the eye-tracker lens adjusted to 60 cm. Participants were instructed to limit blinking to the ITI and to maintain gaze at the central fixation dot throughout testing. The spatial position at which Gabor patches appeared in each trial was quasi-random, such that within every grouping of 15 trials a Gabor would be shown at each of the five spatial positions exactly three times. To assess awareness of the US/CS+ association, after every 15 trials (3 US/CS+ pairings) in the acquisition block participants used a mouse cursor to indicate the screen location where they expected the Gabor to be located when the US noise was played.

### Data Recording and Analysis

#### Eye tracker

Pupil and gaze were recorded continuously at 500 Hz with an EyeLink 1000 Plus (SR Research) eye tracker using a 16-mm lens. Pupil diameter was approximated via ellipse mode, and the illumination level of the infrared signal was set to 100%. To calibrate and validate the eye-tracker, participants fixated on a nine-point grid, which showed a white circle (5 degrees visual angle) at nine locations against a black background. Pupil segments were discarded if the diameter was not between 0.01 and 5 mm (approximately 400–16,000 in default manufacturer arbitrary units). Offline, pupil trials were rejected if artifacts such as blinks obscured more than 200 sample points in an epoch, or if they contained any eye movements. Brief (< 200 sample points) disruptions in the data were interpolated using piecewise cubic interpolation (Mathôt, 2013). To evaluate compliance with fixation instructions, we calculated the proportion of time during which the eye gaze was outside a 2-degree area centered around the fixation cross. This was done separately by experimental condition, only for the time period where experimental stimuli were on the screen. A simple ratio of time outside fixation divided by the presentation time was used.

#### EEG

EEG was recorded continuously from a 128 channel Electrical Geodesics (EGI) system at a sampling rate of 500 Hz, with the vertex electrode (Cz) as recording reference. Data recording was constrained by Butterworth filters both online (low-pass 3-dB point at 170 Hz, high-pass 3-dB point at 0.05 Hz) and off-line (low-pass 40 Hz, 18th order, 3-dB point at 45 Hz, high-pass 4 Hz, second order, 3-dB point at 18 Hz). After arithmetically transforming the data to the average reference, artifact rejection was performed according to the SCADS (statistical correction of artifacts in dense array studies) procedure described in Junghöfer, Elbert, Tucker, and Rockstroh (2000). Electrode impedances were kept at <50 kΩ, as recommended for EGI high-impedance amplifiers.

##### Source Localization and Sensor Selection

Estimated sources of neuronal activity underlying the scalp-recorded EEG were estimated using the classical minimum norm method as described in Hauk, 2004. Using a three-shell source reconstruction model, artifact free epochs of voltage data (performed separately for alpha-band and ssVEP signals) were averaged for each spatial position and source localized, with the 129 sources (out of 655 total estimated model sources) located closest to the electrode positions serving as the time-varying measure of each respective EEG signal. Alpha-band and 15 Hz ssVEP signals were isolated as described below for each of the three orthogonal orientations of the MNE (1 radial, two tangential relative to the scalp surface). For each signal, the three dipole orientations at each source location were then combined by means of the modulus (Euclidean distance) to produce a measure of current density (nanoamperes/mm^2^).

Current density measures at the 129 electrode locations then entered a spatial PCA to determine the relative sensor weighting to apply to each of the five spatial positions of visual stimulus presentation. The PCA was performed (MATLAB function ***pca***) separately for the current densities of the alpha-band time series and the FFT-derived ssVEP amplitude (see below). The average of full trial (2000 ms) post-stimulus onset alpha timeseries data and the ssVEP FFT amplitude from the second (1000 ms) half of each trial following stimulus onset, averaged for each participant at each of the five spatial position conditions in the habituation trial block, entered the PCA. These time ranges were determined based on a body of previous work showing that alpha-power changes in response to conditioned stimuli occur early and extend over 1-2-seconds, whereas ssVEP differences occur predominantly in the second prior to US onset (Wieser, Miskovic, & Keil, 2016). Only habituation trials were used for determining topographical sensor weighting values to avoid possible topographical biases or artifacts induced during acquisition. The first principal component of the PCA of each spatial location (figure 3) was taken as a sensor weighting vector and projected back onto each respective signal, with the resulting measure representing the PCA-weighted average of source localized current density activity for each participant viewing each location (at each timepoint for the alpha timeseries). An alternative time window (not shown) of maximum alpha decrease (300-900 ms, determined by inspection of baseline-divided grand mean power in the habituation trial block) was also examined. While displaying a more retinotopic pattern of activity, results did not differ from using the sensor weights (see below) derived from the first principal component obtained using the full-trial average described above.

**Figure 3.**
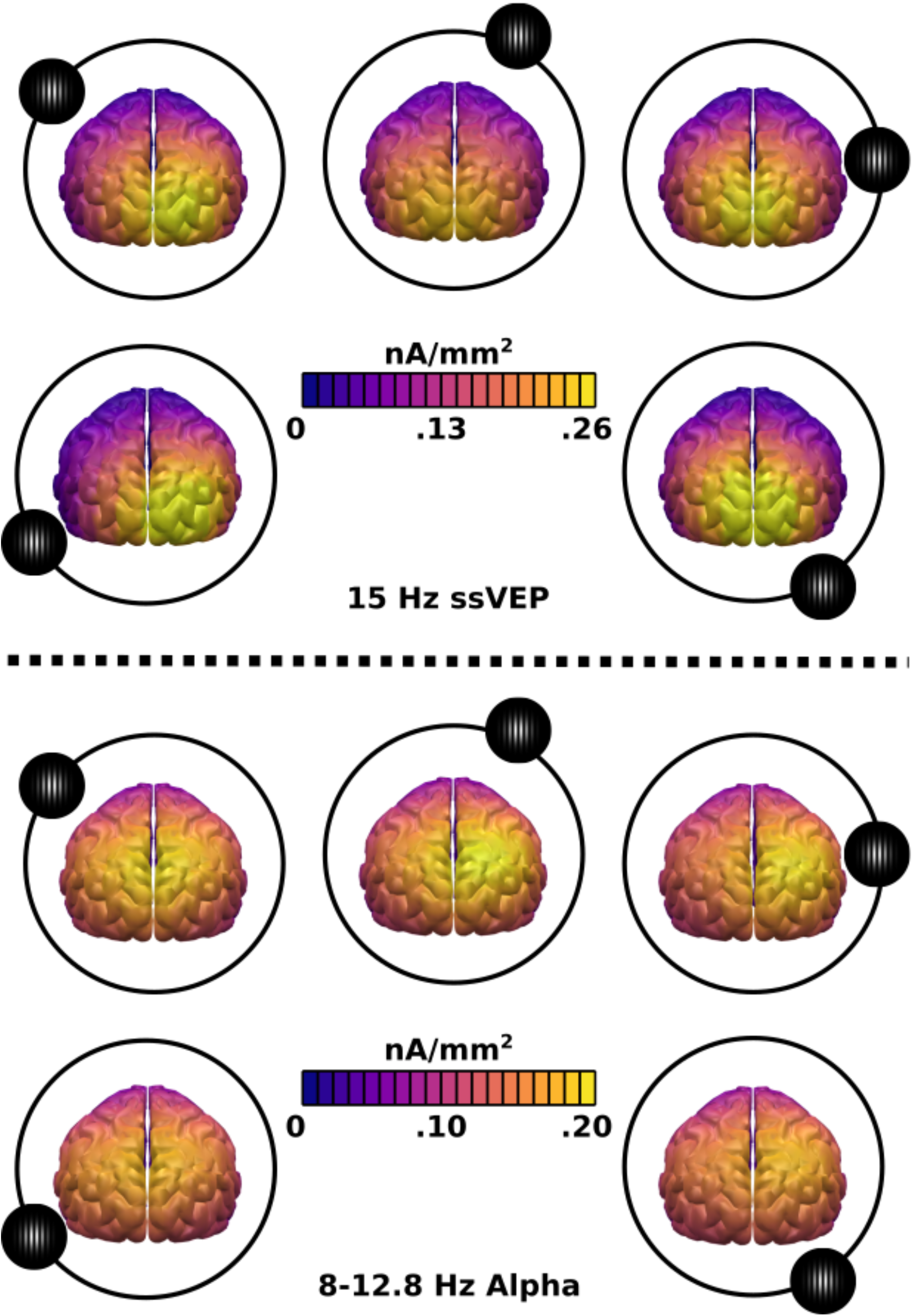
PCA topographies. Topographical activation patterns show the first principal component of source-localized responses to Gabor stimuli appearing at each depicted location during habituation trials.

###### Alpha-band Power

Alpha-band activity was estimated using Morlet wavelet convolution (Tallon-Baudry & Bertrand, 1999; Bertrand & Pantev, 1994) with a Morlet constant (analysis frequency divided by the width of the wavelet in the frequency domain) of 7 to balance resolution of alpha activity in the time and frequency domains. Wavelets with a center frequency between 8 and 12.8 Hz were used to quantify alpha-band changes.

###### Steady-state Amplitude and Signal-to-noise Ratio

The amplitude of the 15 Hz ssVEP response was calculated by means of a moving window fast Fourier transformation (FFT) for a 1000 ms. interval ranging from 1000 – 2000 ms. post stimulus onset. The signal-to-noise ratio (SNR), a measure of the degree to which the strength of a signal of interest exceeds that of surrounding signal “noise” (Teplan, 2002), was calculated as the ratio of the 15 Hz frequency bin to the average of a range of 3 frequency bins above and below 15 Hz (excluding the frequency bins immediately preceding and following 15 Hz; figure 4).

**Figure 4.**
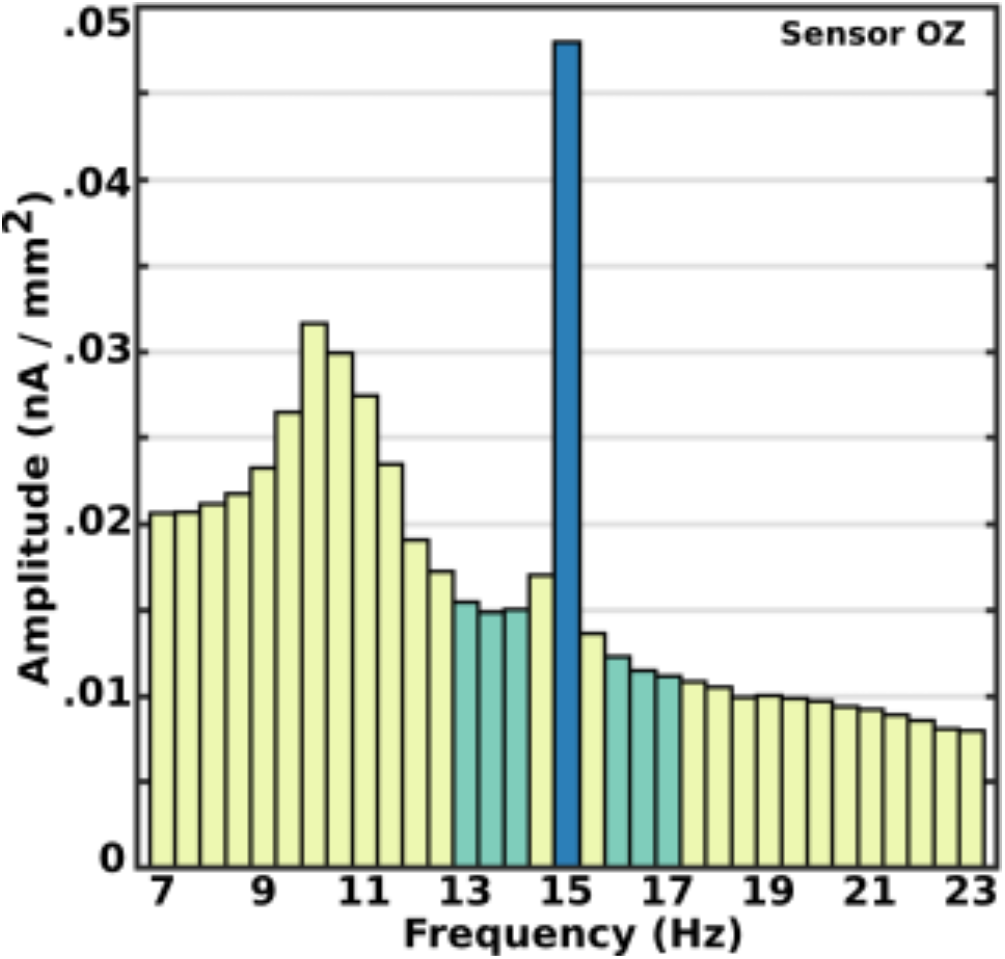
Frequency spectrum of ssVEP. The grand mean of source-localized 15 Hz ssVEP amplitude at one central occipital sensor location, demonstrating that steady-state responses were tightly confined to the driven frequency. Frequency bins used to compute the signal-to-noise ratio (13-14 Hz and 16-17 Hz) are shown in green.

### Statistical Analysis

All statistical inference was performed with Bayes factors (BFs; Jeffreys, 1998; Wetzels et al., 2011) using JZS (Jeffreys–Zellner–Siow) priors (Rouder et al., 2009) and scale factors of 0.707 for two group comparisons. We utilized Bayesian hypothesis testing given the problems of using *p*-values specifically (e.g. Edwards, Lindman, & Savage, 1963; Wagenmakers, 2007) and dichotomous decision criteria generally (McShane, Gal, Gelman, Robert, & Tackett, 2019) to summarize research results. Bayes factors intrinsically control for type-I errors (Gelman and Tuerlinckx, 2000; Gelman, Hill, & Yajima., 2012; Johnson, 2013) and require no adjustments for multiple comparisons, stopping rules, or *a priori* versus *post hoc* comparisons (Dienes, 2008, 2014). Additionally, our specific hypotheses as to the direction and trend of signal change for alpha-band and ssVEP activity were pre-registered (https://osf.io/h7tqa/?view_only=eee3c81e66854d1bb1723ea25cd07f5d). Trials for each of the five spatial locations were averaged across trials within participants, separately for habituation and acquisition trial blocks, using epochs of 3000 msec (500 msec pre- to 2500 msec post stimulus onset) for ssVEP and alpha and 4000 ms (1000 ms pre- to 3000 msec post stimulus onset) for pupil data. As the hypotheses of interest concerned the effect of relative distance from an aversively-paired spatial location, not differential responses at these locations themselves (due, for instance, to differences in the volume of cortical tissue sensitive to different regions of the visual field), all EEG analyses rotated the signal data from viewing each spatial position such that the five spatial positions were transformed into five bins representing relative distance from the conditioned spatial location. In other words, absolute spatial position was discarded while position as a function of distance from the CS+ was conserved and aligned across all participants.

#### Topographical model comparison

Signed *F*-contrasts (Rosenthal, Rosnow, & Rubin, 2000; Rosnow, Rosenthal, & Rubin, 2000), weighting each spatial position to account for the hypothesized trend (gaussian or difference-of-gaussian) of the change in each signal following aversive conditioning, were calculated across the entire source-localized scalp topography. At each source location assigned to the original 129 sensor sites, *F*-values quantified how closely the signal change across spatial positions followed each of the hypothesized trends at that location. The square root of these *F*-values (Rosenthal, Rosnow, & Rubin, 2000, p. 41) were then converted to JZS Bayes factors as detailed by Rouder and colleagues (2009).

#### Effect of aversive conditioning

Dependent variables of interest (ssVEP, alpha, and pupil diameter) were analyzed separately. The 129 channel EEG data for each spatial position was multiplied by its corresponding PCA activity value to produce a weighted 1-dimensional signal value. The two spatial distance bins either one or two position steps away from the CS+ location were merged, with all subsequent analyses performed on the resulting three bins of data (CS+, near, far). Cohen’s *d* was calculated for each signal for each of these 3 location conditions as a standardized measure of the effect of aversive conditioning. For alpha and pupil data timeseries, a JZS Bayes factor (Rouder et al., 2009) was computed for each location condition at each timepoint to quantify the likelihood that these signals differed in acquisition as compared to habituation trials. As time information is lost in the FFT procedure used in the ssVEP analysis, a 95% highest density (credible) interval (HDI) was instead calculated to evaluate the hypothesis of learning-induced change in the ssVEP at each location condition. The median and 95 % HDI of ssVEP SNRs were estimated using ***lmBF*** in the **bayesfactor** package (Morey & Rouder, 2018) for R (R Core Team, 2018), with distance from the CS+ location as the IV and a difference score (SNR in acquisition – SNR in habituation) for each participant as the DV. After estimating a posterior distribution, median and 95% HDIs were calculated on a random sample (10,000 draws) of difference score values from the posterior.

### Code Accessibility

Code and preprocessed, minimum-norm localized (whole-trial averaged for alpha band activity) data for performing the main model comparisons can be freely obtained at this project’s OSF page (DOI 10.17605/OSF.IO/H7TQA).

## Results

### Alpha-band power

Mean alpha band power during habituation was found not to differ between the five spatial positions of Gabor presentation (BF_01_ = 92.97, i.e. favoring the null hypothesis of no difference as a function of location about 93:1 to the alternative hypothesis). The relative equivalence of alpha responses across spatial positions prior to conditioning supports the notion that observed changes in the magnitude and trend across visual field locations result from the aversive conditioning experience.

Fits for the gaussian and DoG models of mean power change from habituation to acquisition across posterior locations are shown in figure 5. Neither tested pattern of mean power change described the observed pattern of selective alpha power decrease well. Instead, both the overall timecourse (figure 6) and the timecourse of the conditioning effect (acquisition trials versus habituation trials; figure 7) suggests a model of binary CS selectivity (CS+ > all non-CS).

**Figure 5.**
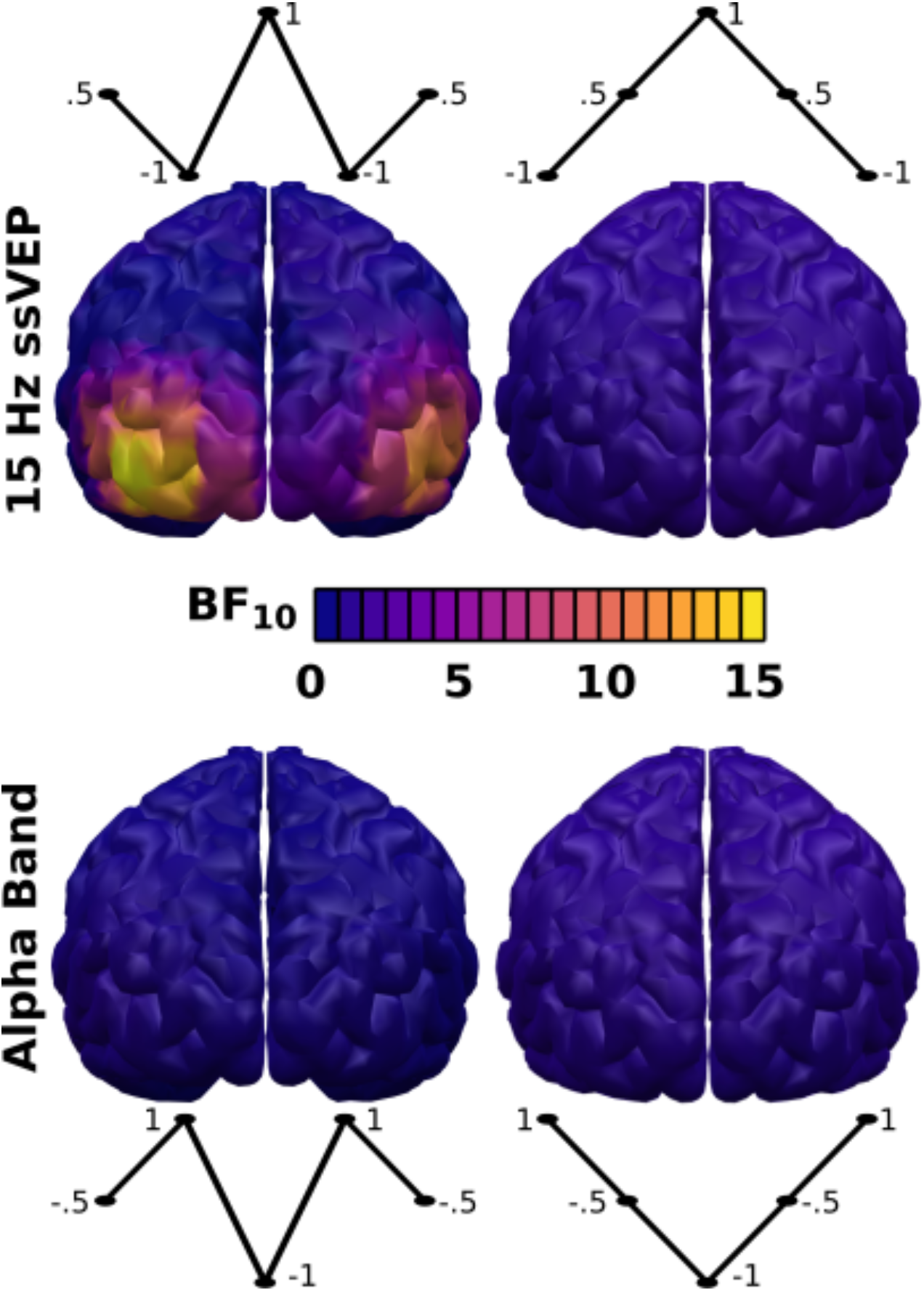
Topographical fits of competing models. Difference-of-gaussian (left) and Gaussian (right) models of source-localized signal change following conditioning is shown for posterior topographical locations for steady-state (top) and alpha band (bottom) signals.

**Figure 6.**
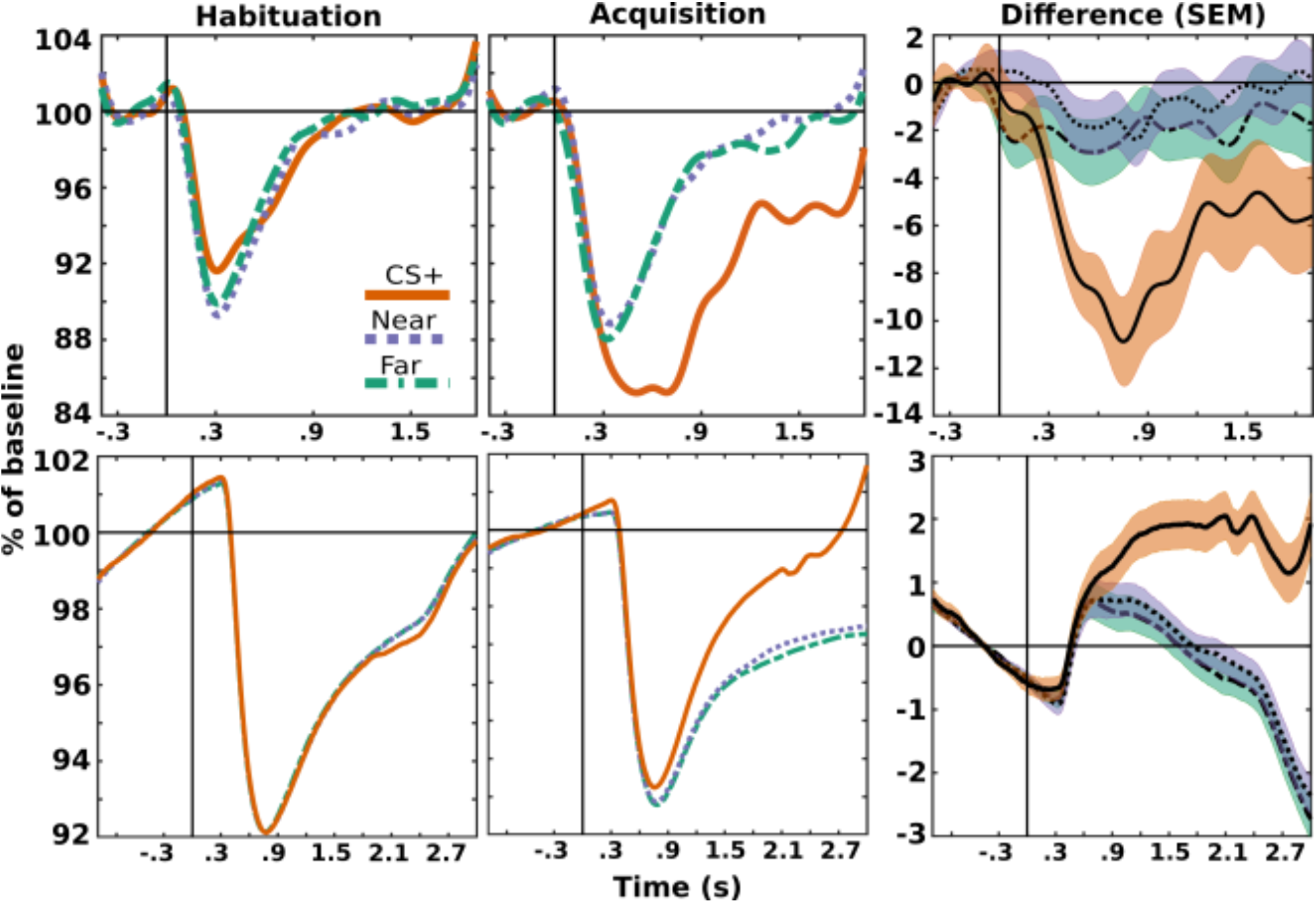
Alpha-band and pupil response time course. Alpha (top) is shown as a percent of mean baseline (400 - 100 milliseconds pre-stimulus onset) power. Pupil response (bottom) is shown as a percent of mean manufacturer (SR Research) reported arbitrary units (1000 – 100 milliseconds pre-stimulus onset).

**Figure 7.**
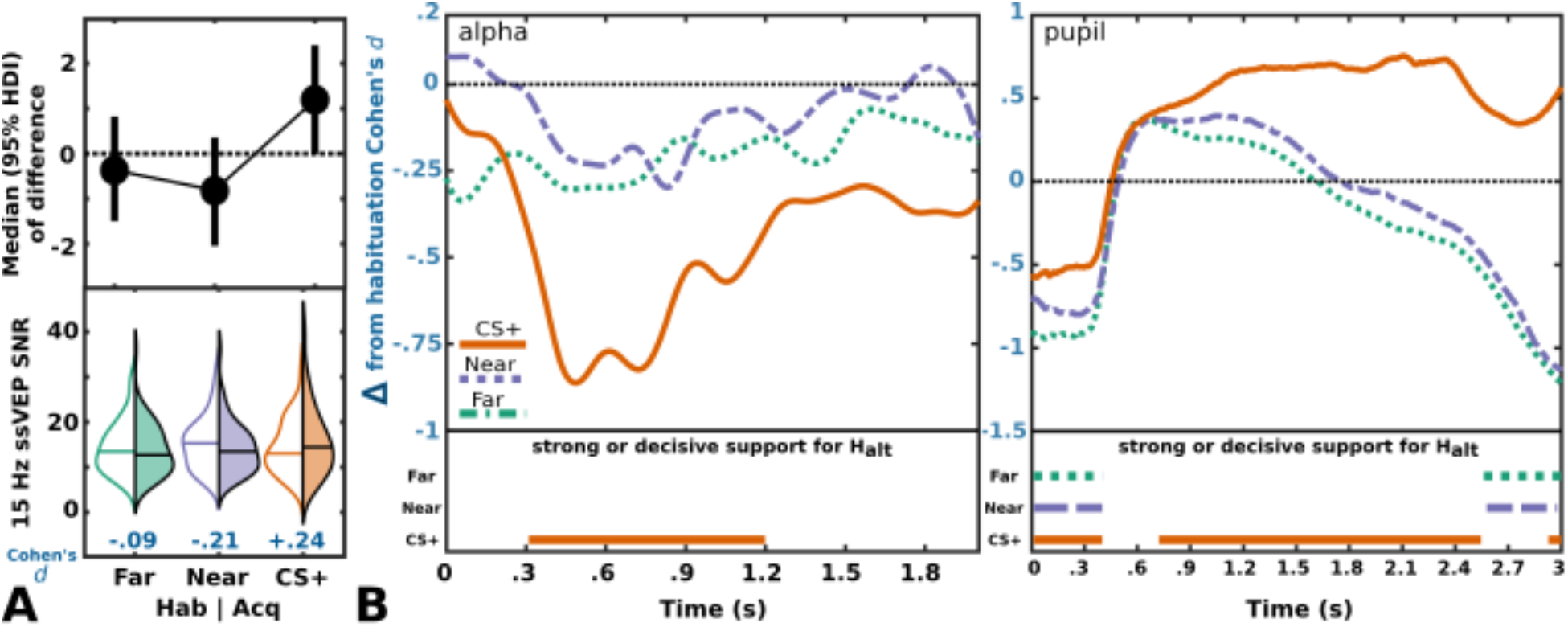
Conditioning effects. Magnitude of change from habituation to acquisition trials as a function of spatial distance from an aversively paired location. A) Distribution (bottom) of PCA-weighted SNR values for all participants in habituation (left violin) and acquisition (right violin). Top plot shows the median and 95 % HDI of difference scores (acquisition – habitation). B) Levels of support generally follow the common heuristics suggested by Jeffreys (1961) for Bayes factors: 3-9 is anecdotal, 10-99 is strong, 100+ is decisive. Only timepoints where support was strong-to-decisive in favor of the alternative hypothesis are indicated in the bottom portion of each figure. Cohen’s *d* effect size of PCA-weighted alpha band activity (as a percentage of mean activity 400 - 100 milliseconds pre-stimulus onset) for the entire post-stimulus timecourse (left). Cohen’s *d* effect size of pupil activity (as a percentage of mean activity 1000 - 100 milliseconds pre-stimulus onset) for the entire post-stimulus timecourse (right).

### Steady-state power signal-to-noise ratio

During habituation, the 15 Hz steady-state SNR was found not to differ between the five spatial positions of Gabor presentation (BF_01_ = 11.62), again suggesting that observed conditioning effects do not stem from pre-existing differences.

Fits for the gaussian and DoG models of mean power change from habituation across posterior and occipital topographical locations is shown in figure 5. There was strong evidence (Jeffreys, 1961) that activity within regions consistent with retinotopic stimulus processing was altered in a manner well described by the DoG model. The PCA-weighted conditioning effect (acquisition trials versus habituation trials; figure 7) largely mirrored the overall topographical trend, with SNR increased for acquisition relative to habituation trials at the CS+ location, suppressed at near unconditioned locations, and relatively unchanged at far unconditioned locations.

### Pupil and gaze

Proportion of deviations from fixation did not vary by spatial position during habituation trials (BF_01_ = 96.48), implying that no stimulus presentation position differentially impacted participants’ ability to maintain fixation. More importantly, the change in proportion of fixation deviations from habitation to acquisition did not vary by aligned location condition (i.e. it did not differ for the CS+ compared to other locations; BF_01_ = 91.9), ruling out the possibility that heightened neural responses to CS+ locations were caused by preferentially foveating them. Similar to the alpha band results presented above, both the overall timecourse (figure 6) and the timecourse of the conditioning effect (acquisition trials versus habituation trials; figure 7) indicate a model of binary CS selectivity (CS+ > all non-CS).

### Awareness of CS+/US association

Overall, approximately 90% (46/51) of participants consistently identified the spatial relationship between the US and the CS+ correctly, indicating that results reported here are unlikely to have been impacted by ambiguity regarding the associative contingency. The time course of mean and individual participant expectancy reports across acquisition trials is shown in figure 8.

**Figure 8.**
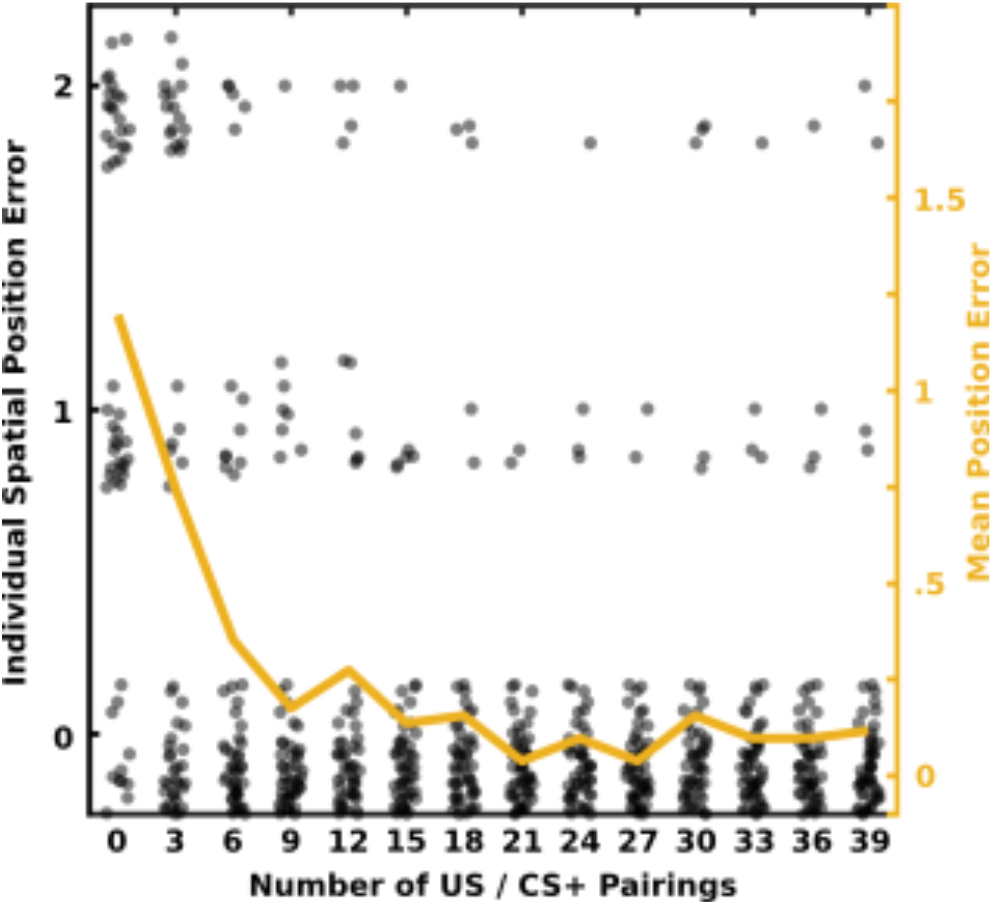
Conditioning contingency awareness. Awareness of the association between the CS+ location and the US noise is shown for each individual (black dots) and for the average over participants (orange line) during the acquisition trial block. The first assessment (x=0) was done at the beginning of the acquisition trial block prior to any US presentations (i.e. participants were randomly guessing the position at which they would hear the US). Random jitter was added to individual responses for purposes of illustration.

## Discussion

The present study examined experience-dependent changes in visuospatial representation wrought by a brief (~30 min) aversive conditioning regimen in which one on-screen location was paired with an aversive noise stimulus while leaving four equidistant locations unassociated. Comparing pupil diameter, alpha-power reduction, and ssVEP enhancement between acquisition and habituation, we found evidence for experience-related changes in all measures. The effect of conditioning on stimulus-induced alpha power reduction was tightly confined to the CS+ condition, displaying no tendency to generalize to either nearby or more spatially distant, non-paired locations. Pupil activity, an index of autonomic engagement with the stimulus, largely mirrored the CS+ stimulus specificity and overall timecourse of alpha band responses (figure 6), providing converging evidence of experience-dependent changes across different functional systems. In contrast to the conditioning effects observed in alpha power and pupillary responses, visuocortical ssVEPs followed a difference-of-gaussian pattern. Consistent with sharpened tuning of visuospatial representations in visual cortex, we observed an experience-driven reduction of cortical responsiveness to proximal unpaired screen locations accompanied by relative enhancement of responses to the CS+ location.

Phasic reduction in alpha-band power following presentation of a visual stimulus has been broadly linked to numerous cognitive functions, most notably arousal and attention. Several theoretical models, including arousal-biased competition (ABC; Mather & Sutherland, 2011) and motivated attention (Lang, Bradley, & Cuthbert, 1997), link processes of arousal to concepts of attention and resource competition. While an overall elevation of general arousal levels is expected during conditioning trials (Lissek et al., 2008), the specificity of the alpha-band response to conditioned stimuli observed here indicates that this effect is more consistent with focused enhancement brought about through attentive processing (Posner & Petersen, 1990). Like reduced alpha-band responses, pupil dilation has long been positively associated with arousal level (e.g. Loewenfeld, 1958), but more recently has also been linked to the employment of covert attention (Naber, Alvarez, & Nakayama, 2013; Olmos-Solis, van Loon, & Olivers, 2018; reviewed in Mathôt, 2018). Because the present study did not aim to disentangle the often co-active processes of attention and arousal, it is plausible that both sources of bias contribute to the selective facilitation of stimuli presented at CS+ locations.

Posner’s (1980) seminal work established that covert, spatially directed attention enhances processing at attended locations. Such shifts are typically prompted by explicit attention cues, conveyed through verbal or on-screen instructions (Sprague & Serences, 2013; Buschman & Kastner, 2015). Here, we show that an experientially acquired spatial association with an unpleasant stimulus prompts a similar facilitation of neural mass activity in visual cortex for stimuli presented at a conditioned location. Conditioning studies manipulating features other than visual field location (e.g., McTeague et al., 2015; Friedl & Keil, 2020) provide evidence that such facilitation manifests differently for alpha band and ssVEP signals.

The same all-or-none alpha-power modulation following aversive conditioning emerged in a previous study where centrally presented Gabor patches varied along a narrowly spaced continuum of spatial frequencies (Friedl & Keil, 2020). This is notable, as it suggests that alpha change circumscribed to a conditioned stimulus does not depend upon the conditioned stimulus being as perceptually distinct as were the easily discriminated spatial positions employed in the current paradigm. Results presented here indicate that selective alpha power reduction and pupil diameter both index binary discrimination between threat and non-threat cues. Together with evidence that the spatial resolution of focused visual attention is substantially inferior to that of visual perception (He, Cavanagh, & Intriligator, 1996; Intriligator & Cavanagh, 2001), modulation of responses to surrounding non-conditioned stimuli appears to require an additional explanatory mechanism.

The present study used ssVEPs to test the hypothesis that retinotopic visual cortex possesses mechanisms for enhancing the spatial contrast between representation of threat-associated versus non-associated locations. Specifically, we considered the role of suppressive interactions between cortical representations of nearby spatial locations as a means of sharpening macroscopic cortical tuning to a threat location. Within the visual system, suppression mechanisms are thought to contribute to transforming broadly tuned excitatory outputs from lateral geniculate nucleus (LGN) relay cells into the more narrowly feature selective responses observed in visual cortex (Shapley, Hawken, & Xing, 2007; Isaacson & Scanziani, 2011; Angelucci et al., 2017; but see Priebe & Ferster, 2008). The difference-of-gaussian pattern of visuocortical response changes across the range of tested spatial positions observed here closely mirrors responses to orientated gratings (McTeague, Gruss, & Keil, 2015) and faces (Stegmann, Ahrens, Pauli, Keil, & Wieser, 2020) parametrically varied in similarity to conditioned threat cues. The finding that altered tuning of the ssVEP emerges within visual cortex whether the low-level visual feature predicting an upcoming aversive event is a particular orientation or a specific visual field location points to response sharpening as a common mechanism that is operating at this early stage of the associative learning process. An open question, however, is if such sharpening results exclusively from altered sensitivity in feature-specific neuronal populations (e.g., Summerfield & Egner, 2017; Liu, 2019), or if a learned categorical distinction (threat vs. safe) also contributes (Dunsmoor, Kragel, Martin, & LaBar, 2014).

Narrowly tuned response profiles, as observed in ssVEPs here, diverge from what would be expected under generalization models based purely upon perceptual discrimination (e.g. Lissek, 2012; Lissek et al., 2014). Following a perceptual discrimination model, response strength should decrease monotonically as stimuli become more distinct from the CS+ (Onat & Buchel, 2015). Onat and Buchel (2015) instead cite the active integration of “hyper-sharp” responses within several brain regions including the hippocampus and anterior insula with more broadly tuned responses from other regions as responsible for producing the stereotypical gaussian-shaped generalization gradient. While Onat and Buchel (2015) assert that generalization is not solely driven by perceptual ability, Tuominen et al. (2019) found that measures of skin conductance, expectancy ratings, and fMRI BOLD responses in anterior insula and superior frontal gyrus only showed signs of generalization when discrimination performance was low (i.e. near individual perceptual thresholds). Both the significance of sharply tuned neural response profiles such as the difference-of-gaussian to perceptual discrimination, as well as the relationship between response gradients measured at the behavioral and neural level, have strong translational and clinical implications (Dunsmoor & Paz, 2015) and remain pressing questions for future study.

The generally well-defined structural and functional properties of the visual cortex make it an appealing model system for investigating fundamental brain processes (Creutzfeldt, 1977; Priebe & Ferster, 2008) such as associative learning. For nearly a century, behavioral response gradients have provided insight into how the properties of a conditioned stimulus transfer to similar but non-conditioned stimuli. The present work extends the use of this analytical method to the neural population-level, examining the extent to which conditioning an on-screen stimulus location influences visuocortical responses to objects appearing at surrounding visual field locations. Illustrating the dissociable signatures of early visuocortical (ssVEP) and alpha-band responses to spatially contingent aversive events highlights two mechanisms employed by the visual system in adapting to dynamic environments.

## Acknowledgments

This research was supported by grants from the National Institutes of Health (R01MH112558 and R01MH097320) and Grant N00014-18-1-2306 from the Office of Naval Research to A. K.

